# Effects of language experience on domain-general perceptual strategies

**DOI:** 10.1101/2020.01.02.892943

**Authors:** Kyle Jasmin, Hui Sun, Adam T. Tierney

**Author notes:** Correspondence to: Kyle Jasmin. equal contributions (Kyle Jasmin, Hui Sun).

## Abstract

Speech and music are highly redundant communication systems, with multiple acoustic cues signaling the existence of perceptual categories. This redundancy makes these systems robust to the influence of noise, but necessitates the development of perceptual strategies: listeners need to decide the importance to place on each source of information. Prior empirical work and modeling has suggested that cue weights primarily reflect within-task statistical learning, as listeners assess the reliability with which different acoustic dimensions signal a category and modify their weights accordingly. Here we present evidence that perceptual experience can lead to changes in cue weighting which extend across tasks and across domains, suggesting that perceptual strategies reflect both global biases and local (i.e. task-specific) learning. In two experiments, native speakers of Mandarin (N=45)—where pitch is a crucial cue to word identity—placed more importance on pitch and less importance on other dimensions compared to native speakers of non-tonal languages English (N=45) and Spanish (N=27), during the perception of both second language speech and musical beats. In a third experiment, we further show that Mandarin speakers are better able to attend to pitch and ignore irrelevant variation in other dimensions compared to English and Spanish speakers, and even struggle to ignore pitch when asked to attend to other dimensions. Thus, an individual’s idiosyncratic auditory perceptual strategies reflect a complex mixture of congenital predispositions, task-specific learning, and biases instilled by extensive experience in making use of important dimensions in their native language.

## 1. General Introduction

Speech and music are highly redundant communication systems: any given structural feature tends to be conveyed by multiple acoustic cues, some of which are more informative than others. For example, there are many cues to voicing (the phonetic feature that distinguishes, e.g., /b/ from /p/), such as the pitch of the following vowel, but the most important cue relates to the timing of the onset of vocal fold vibration (Voice Onset Time; Massaro and Cohen, 1977; Lisker, 1986). At the level of prosody (the fluctuations of pitch, duration and amplitude that span multiple syllables), there are multiple cues to the presence of features such as stress, which, in spoken English, is associated with pitch movements, longer duration, increased amplitude, and changes to the spectral characteristics of the vowel (Fear, Cutler & Butterfield, 1995; Plag, Kunter & Schramm, 2011). In music, accents are associated with longer note duration and sudden changes in pitch (Ellis and Jones, 2009; Prince, 2014). This redundancy - multiple cues providing the same information - is a highly useful feature because it enables speech and music to be robust to distortion of auditory cues, either due to the presence of masking noise (Winter, 2014) or to noise added by the intrinsic variability of an individual’s perceptual system (Jasmin, Dick, Holt & Tierney, 2020a). (If a cue cannot be perceived, another one can be used instead). However, redundancy also poses a problem for listeners, who need to decide how to integrate multiple sources of sometimes conflicting information.

How do listeners converge on perceptual strategies, i.e. tendencies for certain dimensions to be ‘weighted’ more than others during perceptual categorization? Recent computational models of the development of dimensional weighting in a listener’s first language suggest that listeners prioritize dimensions which have been informative in the past, i.e. dimensions in which the distribution of values form two or more clear peaks which are clearly separated, such as Voice Onset Times for voiced and voiceless consonants (Toscano and McMurray, 2010). More generally, perceptual experience may be able to modify dimensional weighting, as dimensions which tend to be task-relevant are favored over less commonly relevant dimensions. Here we ask whether effects of perceptual experience on dimensional weighting are strictly limited to a particular categorization task in a particular domain, or whether perceptual experience can lead to more sweeping cross-domain changes in dimensional weighting.

One way to test the idea that differences in perceptual experience can modify dimensional weighting is to examine the effects of experience learning different languages. Acoustic dimensions play different roles across languages. In Mandarin, for example, each syllable is pronounced with a ‘tone’ (high, rising, falling-rising, and falling), a pitch contour that is an integral part of the word and helps determine its meaning. Thus, two words with identical speech sounds (consonants and vowels) can have two entirely different meanings depending on the fundamental frequency patterns of the voice used when saying them. For instance, *ma1* means ‘mother’ while *ma3* means ‘horse’ (with numbers indicating tone patterns). In non-tonal languages such as English, on the other hand, pitch has a much smaller influence on word identification (Idemaru & Holt 2011), and instead plays a more secondary role. It can, for example, convey phrase structure and emphasis, features that are also redundantly conveyed by other dimensions, such as duration and amplitude (Streeter, 1978; Mattys, 2000; Chrabaszcz, Winn, Lin & Idsardi, 2014).

The cross-linguistic differences in the role of auditory dimensions in signaling linguistic categories have led researchers to ask whether language experience can affect the strategies listeners use when integrating different sources of evidence during categorization of speech sounds. Native speakers of Mandarin, for example, have been shown to rely more heavily on pitch than native English speakers when they categorize and produce English stress (Nguyễn, Ingram & Pensalfini, 2008; Wang, 2008; Zhang, Nissen & Francis, 2008; Yu and Andruski, 2010; Zhang and Francis, 2010; but see Chrabaszcz et al., 2014) and phrase boundaries (Zhang, 2012). These findings suggest that dimensions which provide particularly important cues to speech categories in a listener’s native language (L1) are up-weighted during second language (L2) perceptual categorization, while dimensions which provide less informative cues are down-weighted, leading to perceptual strategies which differ from those of native speakers. The influence of language experience on perceptual strategies has also been demonstrated for segmental speech perception; for example, native Japanese speakers down-weight F3 and up-weight F2 when perceiving /l/ versus /r/ in English (Iverson et al. 2003).

Two types of models have been posited to account for effects of language experience on speech perception strategies. Perceptual interference models of second language speech perception (Iverson and Kuhl, 1994; Flege, 1995; Best, McRoberts & Goodell, 2001) posit that second language speech categories are perceived via reference to existing language speech categories. For example, according to these models, when native Mandarin speakers categorize English syllables as stressed or unstressed, they refer to tone categories in Mandarin, and so place more weight on pitch information (and less on other cues) compared to native English speakers, given that pitch is by far the most important dimension for distinguishing between lexical tones (although other cues such as duration can be used if pitch is neutralized, such as in whispered speech; Lin and Repp, 1989; Blicher, Diehl & Cohen, 1990; Whalen and Xu, 1992; Fu and Zeng, 2000; Liu and Samuel, 2004). Attentional theories of how information from different acoustic dimensions is combined during speech perception suggest that dimensions that have a greater tendency to capture attention are more strongly weighted (Gordon, Eberhardt & Rueckl, 1993; Francis and Nusbaum, 2002; Holt et al., 2018). According to these models, repeated task-relevance of a particular dimension leads to an increase in perceptual salience of that dimension, thereby increasing the weight that is assigned to that dimension as a source of evidence during perception.

Because these models were designed to account for linguistic data, none of them make claims about whether native experience with tonal language might affect cue weights outside of speech, domain-generally. Indeed, no prior work has investigated whether language experience can modify non-verbal dimensional weighting, i.e. the strategies used when information is combined across multiple dimensions during sound perception. Categorization of sounds across many different domains requires integration of pitch with information from other dimensions. For example, phrase boundaries in music are characterized by changes in both pitch (a shift from low to high or high to low) and duration (a shift toward longer notes; Palmer & Krumhansl, 1987; Tierney, Russo & Patel, 2011), musical ‘beats’ are characterized by melodic leaps and duration changes (Hannon, Snyder, Eerola & Krumhansl, 2004; Ellis & Jones, 2009; Prince, 2014), and the material and size of the objects giving rise to impact sounds are conveyed by the frequency and decay rate of partials (Lutfi and Liu, 2007).

Given the partial overlap in the relevant cues between musical perception and speech prosody perception, we hypothesized that effects of language experience on prosodic cue weighting strategies may transfer to music perception. If so, this would suggest that effects of perceptual experience on dimensional weighting are not limited to a particular perceptual task, or even a particular domain, but can extend across domains to potentially affect domain-general baseline perceptual strategies. Testing effects of perceptual experience on domain-general dimensional weighting could also help constrain theories about the neural and cognitive mechanisms underpinning the effects of language experience on cue weighting. For example, if L1 modifies cue weighting domain-generally, this would suggest that these cue weighting shifts reflect modifications to auditory processing that occur relatively early (i.e. in regions of auditory cortex sensitive to domain-general acoustic cues; see Jasmin, Dick, Stewart & Tierney, 2020b), rather than in down-stream domain-specific regions.

Here we investigated perception of English phrase boundaries and musical beats in native speakers of a tonal language (Mandarin Chinese) and two comparison groups—native speakers of English and of Spanish (non-tonal languages). The native Spanish speakers were included as a second comparison group to ensure that differences in English proficiency were not the primary determinant of any differences between the Mandarin and English speakers. In Experiments 1 and 2 we examined dimensional weighting of pitch and duration during perceptual categorization in speech and in music by creating a two-dimensional stimulus space in which the extent to which pitch versus duration patterns implied the existence of a particular structural feature was orthogonally varied. Importantly, this stimulus space contained ambiguous stimuli, for which pitch suggested one interpretation while duration suggested another. There is no “correct” answer regarding how these stimuli should be categorized, and so they are ideal for examining dimensional weighting strategies: when pitch and duration information conflict, participants with strong pitch weighting and participants with strong duration weighting will categorize the stimuli very differently. Participants were presented with each stimulus multiple times throughout the experiment and asked to categorize it as having an early versus late intonational phrase boundary (prosody perception test) or as having duple meter (strong-weak) versus triple meter (strong-weak-weak) (music perception test). We then used logistic regression to calculate cue weights—the extent to which participants’ categorizations were influenced by pitch versus durational information. We predicted that the Mandarin speakers would rely more on pitch and less on other dimensions (i.e. in this case, duration) when perceiving and categorizing music, as well as speech.

Although no prior study has examined the effects of language experience on dimensional weighting during non-verbal sound perception, there is prior evidence suggesting that tone language experience is linked to enhanced pitch sensitivity. More specifically, tone language speakers have been shown to have more precise pitch discrimination (Pfordresher and Brown, 2009; Giuliano et al., 2011; Bidelman, Hutka & Moreno, 2013; Hutka, Bidelman & Moreno, 2015, Zheng and Samuel, 2018; but see Burns and Sampat, 1980; Stagray and Downs, 1993; Bent, Bradlow & Wright, 2006; Peretz, Nguyen & Cummings, 2011), more precise pitch interval discrimination (Pfordresher and Brown, 2009; Hove, Sutherland & Krumhansl, 2010; Giuliano et al., 2011; Creel, Weng, Fu & Heyman, 2018), more precise pitch contour discrimination (Deroche et al., 2019), better melody discrimination (Wong et al., 2012; Bidelman et al., 2013), and better vocal melody production (Pfordresher and Brown, 2009). These results are somewhat orthogonal to our current investigation, as our interest is in the importance or weighting placed on information gained from pitch, rather than the ability to extract information from pitch in the first place. To make sure that our paradigms assessed pitch weighting rather than pitch sensitivity, we measured all participants’ pitch discrimination thresholds and ensured that the differences between pitch levels for stimuli used in our experiment were always greater than the thresholds for all participants.

Our pitch weighting measure was, therefore, theoretically dissociable from pitch sensitivity. Nevertheless, the greater domain-general pitch sensitivity in Mandarin speakers could be one explanation for why they weight pitch more highly during perceptual categorization, as all else being equal, individuals will place greater importance on perceptual channels in which signals are less variable (Ernst and Banks, 2002). Congenital amusics, for example, who have a domain-general deficit in pitch sensitivity, down-weight pitch information during perceptual categorization (Jasmin et al., 2020a). Indeed, Cantonese-English bilingual speakers are better than native English speakers at discriminating English stress based on subtle isolated pitch cues (Choi, Tong & Samuel, 2019), suggesting that up-weighting pitch information might be an optimal strategy for tone language speakers due to their greater pitch sensitivity. We investigated whether group differences in pitch sensitivity are the primary factor driving any group differences in dimensional weighting strategies in two main ways. First, after our initial group analysis of the results from Experiments 1 and 2, we performed a follow-up analysis by matching a subset of the participants for pitch sensitivity and testing whether native language effects on dimensional weighting strategies persisted even after the small group differences in sensitivity were removed. Second, in Experiment 3, we explicitly asked participants to direct attention to the pitch or amplitude of speech while the other dimension orthogonally varied. We also replicated the results of Experiment 3 in our subset of participants that were matched for pitch sensitivity. Our reasoning was that if our results were primarily driven by pitch sensitivity, then we should find no group differences when participants were explicitly asked to attend to amplitude.

## 2. Experiment 1: Native language effects on second language perception

### 2.1. Introduction

Experiment 1 tested whether native experience with Mandarin (a tonal language) leads to development of a perceptual strategy that extends into a second language, English. Participants listened repeatedly to a spoken phrase whose pitch and duration characteristics varied in such a way as to indicate a grammatical phrase boundary earlier or later. Pitch and duration were varied orthogonally, allowing us to measure the degree to which categorization judgments reflected reliance on pitch or duration. Based on previous studies indicating that Mandarin speakers weight pitch more highly when processing English lexical stress (Nguyễn et al. 2008, Wang, 2008, Zhang et al. 2008, Yu and Andruski 2010, Zhang and Francis 2010) and phrase boundaries (Zhang 2012), we predicted that Mandarin speakers would rely more on pitch to perceive English phrase boundaries.

### 2.2. Methods

#### 2.2.1. Participants

Fifty (50) native speakers of British English were recruited from the Prolific online participant recruitment service (prolific.co). An initial automated screening accepted only participants who spoke English as a native language, and an initial questionnaire at the outset of the study confirmed that this was the case by asking them “What is your NATIVE language (i.e. the language FIRST spoken)? If more than one, please list each one.” Fifty (50) native speakers of Mandarin were recruited from an ongoing longitudinal study. The Mandarin speakers all had resided within the United Kingdom for 6-12 months and had not previously lived in an English-speaking country. A second non-tonal language group was recruited, consisting of 30 speakers of Spanish who reside in the UK. The sample size for the Spanish-speaking group was the maximum number of participants that could be recruited and tested given time and resource constraints. All participants gave informed consent and ethical approval was obtained from the ethics committee of the Department of Psychological Sciences at Birkbeck, University of London.

We used the Gorilla Experiment Builder (www.gorilla.sc) to create and host our experiment (Anwyl-Irvine, Massonnié, Flitton, Kirkham & Evershed, 2020). Participants were asked to wear headphones, and automated procedures ensured that participants were all using the Google Chrome browser on a desktop computer. One drawback of online testing is that it can be somewhat more difficult to ensure that participants are fully engaged with the task. In an attempt to minimize spurious data points we only included data from participants for whom, in both Experiment 1 and Experiment 2, there was a significant relationship (*p* < 0.05) between at least one of the stimulus dimensions (pitch or duration) and categorization responses. (See the task descriptions below for more details. This participant inclusion criterion was set a priori, prior to data collection.) This criterion caused the exclusion of five Mandarin-speaking participants, five English-speaking participants and three Spanish speakers, resulting in final group totals of 45 Mandarin speakers (mean age 23.5 ± 1.9, 39 F), 45 English speakers (mean age 25.6±5.2, 21 F), and 27 Spanish speakers (mean age 29.5±6.1, 18 F). All the Mandarin speakers arrived late in a second language environment after the age of 19 (mean age of arrival 22.8±1.8) and had only a short length of residence in the UK (mean years 0.8±0.1). However, they had received an extensive amount of English class training in China (mean years 13.8±2.3). The other non-English group, Spanish speakers, showed greater individual variability in their age of arrival (mean age 26.4±6.1), length of residence in the UK (mean years 3.0±2.0) and English class training (mean years 12.1±4.5). All three groups also reported varied music training backgrounds (i.e., regular training longer than a year). There were 21 Mandarin speakers (mean years 2.9±4.4 for the group), 4 Spanish speakers (mean years 0.4±1.6 for the group) and 16 English speakers (mean years 1.6±3.7 for the group) who reported some music training experience.

#### 2.2.2. Stimuli

First, recordings were made of a Standard Southern British English-speaking voice actor reading aloud two different sentences: “If Barbara gives up, the ship will be plundered” and “If Barbara gives up the ship, it will be plundered”. The first six words of each recording were extracted; these recordings were identical lexically but differed in the placement of a phrase boundary, i.e. after “up” in the first recording (henceforth “early closure”) and after “ship” in the second recording (“late closure”). The speech morphing software STRAIGHT (Kawahara & Irino, 2005) was then used to morph the early closure and late closure recordings onto one another using the standard procedure: the F0 was extracted from voiced segments of the two utterances; next, aperiodic aspects of the signal were identified and analyzed; then, the filter characteristics of the signal were calculated. Finally, the two “morphing substrates” (speech from each recording decomposed into F0, aperiodic aspects and filter characteristics) were manually time aligned by marking corresponding ‘anchor points’ in both recordings, such as the onsets of salient phonemes, so that morphs reflect the temporal characteristics of the two initial recordings, varying in the extent to which acoustic cues imply the existence of a phrase boundary either at the middle or at the end of the phrase. Pitch and duration were set to vary across five morphing levels, expressed as percentages, which included 0% (identical to the acoustic pattern for the early closure recording), 25% (a greater contribution of early closure than late closure recording), 50% (equal contribution from both recordings, and therefore ambiguous with respect to the placement of the phrase boundary), 75% (greater contribution of late closure than early closure), and 100% (identical to the pattern for the late closure recording). In total, therefore, there were 25 stimuli (one stimulus for every unique combination of five pitch and duration levels).

To confirm that the effects of the F0 manipulation were relatively limited to the fundamental frequency and did not have broader effects on spectral shape, we investigated the spectral tilt for each of the five pitch levels, collapsing across all duration levels. Spectral tilt was measured by calculating the long-term average spectrum and then taking the ratio of the energy below 1 kHz and between 1 and 4 kHz (Murphy et al., 2008). Spectral tilts for the stimuli with 0, 25, 50, 75, and 100% pitch level were 4.55, 4.41, 4.31, 4.49, and 4.25, respectively. Therefore, the effects of the F0 manipulation on spectral shape were relatively minimal.

#### 2.2.3. Procedure

Participants read instruction slides and completed practice trials in order to familiarize themselves with the task. During instructions, participants were presented with a clear example of early versus late closure (two full sentences with original, unaltered pitch and duration cues) along with the text and were asked to listen to each example three times before proceeding. During practice trials they were then presented with a short version of these two examples (i.e., the first six words) and asked to categorize it as early or late closure by clicking a button to indicate where the comma is placed. If it sounded more as if the phrase boundary was in the middle, they would click “If Barbara gives up, the ship”; if it sounded more as if the phrase boundary was at the end, they would click “If Barbara gives up the ship,”. After each trial, participants were given feedback as to whether they answered correctly. They were not able to proceed to the main test until they answered both trials correctly. During the test itself, each participant completed 250 trials, ten blocks in each of which the 25 items were presented in random order. Note that, during the actual test, the stimuli were all identical lexically (i.e. they consisted of the same six words) but varied in the extent to which pitch and duration patterns implied the existence of an early or late phrase boundary. The tasks used in Experiments 1, 2, and 3 are available at Gorilla Open Materials (https://gorilla.sc/openmaterials/89766).

#### 2.2.4. Analysis

For each participant logistic regression was conducted to examine the extent to which pitch versus duration influenced their categorization judgments. The outcome variable was the categorization decision for a given trial, with pitch (5 levels) and duration (5 levels) as predictors. The resulting coefficients were then normalized so that they summed to 1 using the following equation:

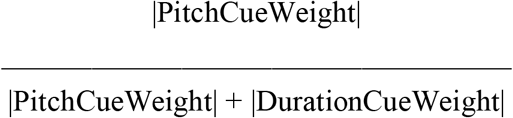

The cue weights were not normally distributed. For this reason, comparisons among all three language groups are reported with the non-parametric Kruskal-Wallis H statistic. Comparisons of two language groups are reported with Mann-Whitney *U* tests. As an effect size measure, we report Vargha-Delaney A, which is the probability that a randomly selected value from one group will be greater than a randomly-selected value from another group.

### 2.3. Results

Normalized cue weights were calculated for each participant. These measures reflect the degree to which participants relied on duration or pitch to make their judgments (see Methods). Cue weights differed across the three language groups (H(2) = 23.35, *p* < .001). For the speech task, native Mandarin speakers had larger normalized pitch cue weights than both native English speakers (U = 1589, *p* < .001, A = 0.78) and native Spanish speakers (U = 854, *p* = .004, A = 0.70), indicating that they relied on pitch to a greater degree and duration to a lesser degree than these groups (Fig. 1a). English and Spanish speakers’ cue weights did not differ (U = 468, *p* = 0.11, A = 0.39). Figures 1b and 1c show how participants’ categorization responses changed as the pitch and duration levels of the stimuli were varied, respectively, collapsing across the other dimension; the categorization function for pitch was steeper for Mandarin speakers than for the other two groups, while the categorization function for duration was shallower for Mandarin speakers than for the other two groups. To investigate which stimuli were driving the group differences in categorization strategies, we used Mann-Whitney *U* tests to compare categorization responses to each of the 25 stimuli, using the Bonferroni method to correct for multiple comparisons. Group differences were almost exclusively confined to ambiguous stimuli in which pitch and duration information suggested conflicting interpretations (Figures 1d and 1e).

**Figure 1.**
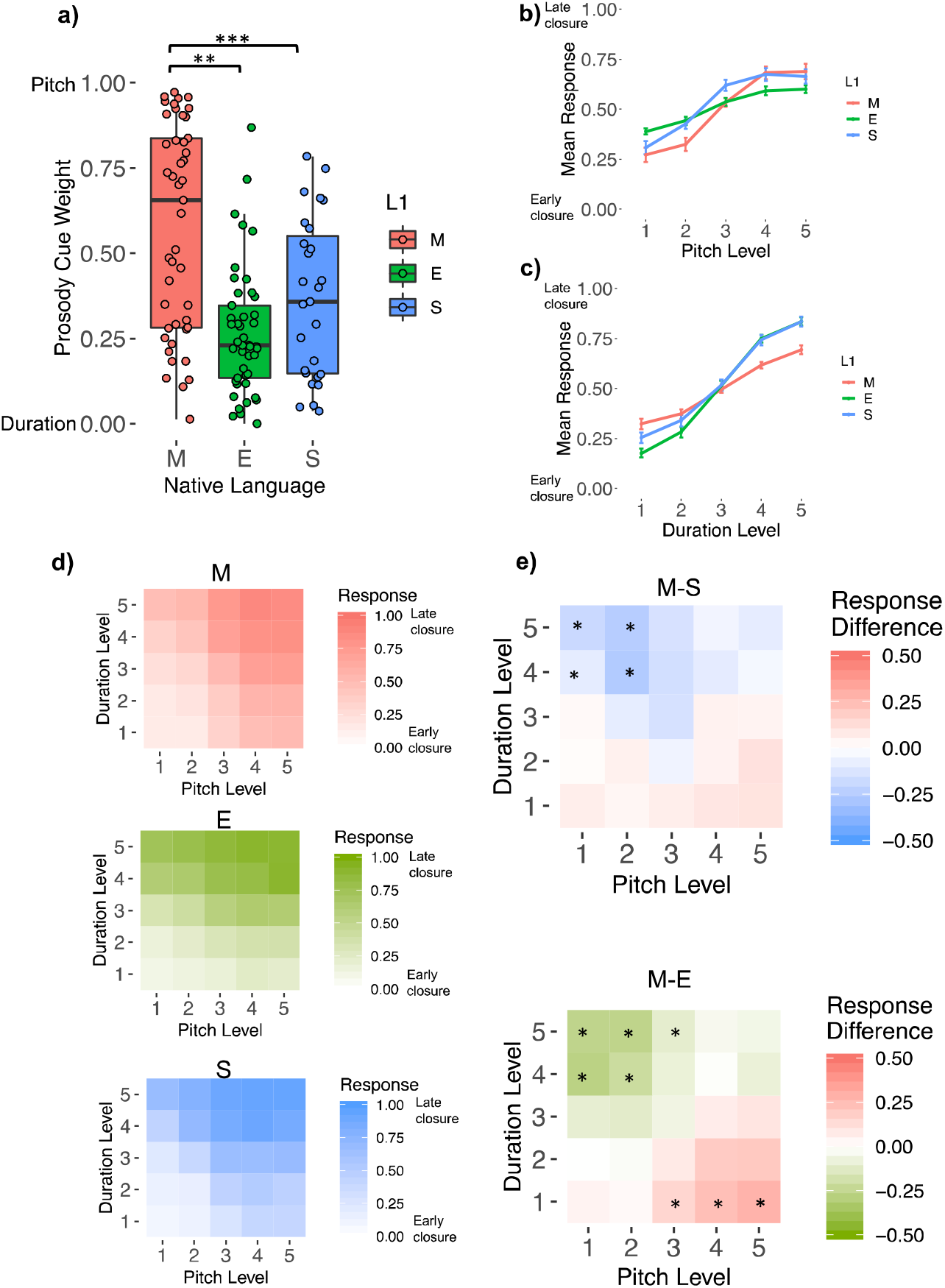
Mandarin speakers rely more on pitch and less on duration when categorizing features in speech compared to English and Spanish speakers. **a)** Plots of normalized cue weights by language group for the speech task. Greater values (approaching 1) indicate greater reliance on pitch, and lower values (approaching 0) reflect greater reliance on duration. **b)** Responses for levels of the pitch dimension (collapsed over duration) and **c)** for the duration dimension (collapsed over pitch). Error bars represent SEM. **d)** Plots of responses during the speech task for each cell of the stimulus space, averaged across participants in each of the groups. Darker colors indicate more “late closure” responses. **e)** Responses from the Mandarin group compared to the English and Spanish groups. Darker colors indicate relatively more “late closure” responses made by the Mandarin (red), or English (green) and Spanish (blue) groups. Asterisks indicate significant group differences in categorization responses, as tested using Mann-Whitney *U* tests with Bonferroni correction for multiple comparisons.

### 2.4. Discussion

As predicted, the native Mandarin speakers relied on pitch information to perceive phrase boundaries in English speech more than both native English and Spanish speakers. The results are in line with previous studies showing Mandarin speakers rely more on pitch and less on other cues to perceive English lexical stress (Nguyễn et al., 2008; Wang, 2008; Zhang et al., 2008; Yu and Andruski, 2010; Zhang and Francis 2010) and phrase boundaries (Zhang, 2012). The comparisons between groups for each of the 25 stimulus cells indicated that categorization behavior differed the most when pitch and duration provided conflicting information. For these cue-incongruent stimuli (top-left and bottom-right corners; Figures 1d and 1e) the Mandarin group tended to base their judgments more on pitch, and less on duration, than the English and Spanish groups. Moreover, group differences were not limited to stimuli with more subtle pitch cues, but extended to stimuli with large, obvious pitch cues, suggesting that the group differences in categorization were not primarily driven by differences in sensitivity. Together these results indicate perceptual strategies that are useful in one’s native language are deployed to understand second languages. In Experiment 2 we investigated whether effects of language experience on perceptual strategies transfer to the perception of music.

## 3. Experiment 2: Native language effects on music perception

### 3.1. Introduction

Experiment 1 demonstrated that perceptual strategies instilled by one’s native language can extend to perception of a second language. The next experiment tested the novel hypothesis that such strategies extend even beyond the domain of language, into perception of other sounds. Musical structure (like linguistic structure) is simultaneously indicated by multiple cues from pitch, duration and amplitude, and therefore provided a suitable testing ground for our hypothesis. We asked participants to listen to musical sequences and judge the locations of musical beats. The stimuli were constructed such that both pitch and duration information cued the locations of beats, orthogonally and to varying degrees. If Mandarin speakers weight pitch information highly during perception of sounds other than speech, their judgments of beat locations should be influenced by pitch information to a greater extent than those of English and Spanish speakers.

### 3.2. Methods

#### 3.2.1. Participants

The same participants took part as in Experiment 1.

#### 3.2.2. Stimuli

In each trial participants heard 18 tones (a group of 6 tones repeated three times) that varied in pitch and duration patterning. Tones were four-harmonic complex tones with equal amplitude across harmonics and a 15-ms cosine ramp at note onset and offset to avoid transients. Pitch and duration patterns each varied across five levels, which differed in the extent to which the cues implied a three-note grouping (STRONG weak weak STRONG weak weak; “waltz time”) versus a two-note grouping (STRONG weak STRONG weak STRONG weak; “march time”). The strength of these groupings was conveyed by varying the pitch of the first note of the 2-note or 3-note groupings relative to the other notes in the grouping. An increase in the pitch of a note implied the existence of a strong beat at that location. Similarly, an increase in the duration of a note implied the existence of a strong beat there.

The five pitch levels were [B A A B A A] (strongly indicating a groups of three), [Bflat A A Bflat A A], [A A A A A A] (no grouping structure indicated by pitch), [Bflat A Bflat A Bflat A], and [B A B A B A] (strongly indicating groups of two), where “A” was equal to A440, i.e. 440 Hz, “Bflat” was equal to 466.2 Hz, and “B” was equal to 493.9 Hz. The duration levels manipulated the duration of notes (not the inter-onset intervals, which were always 250 ms). So, the five duration levels (in ms) were [200 50 50 200 50 50] (strongly indicating groups of three), [100 50 50 100 50 50], [50 50 50 50 50 50] (no grouping conveyed by duration), [100 50 100 50 100 50], and [200 50 200 50 200 50] (strongly indicating groups of two). Crucially, the five pitch levels and 5 duration levels were varied orthogonally, for a total of 25 conditions -- thus pitch and duration sometimes conveyed the same grouping pattern (a group of two or a group of three), and for other stimuli conveyed different, competing patterns. Note also that the size of the cues was kept large enough that they should be detectable by most listeners. Psychophysical thresholds were collected in an attempt to confirm whether our participants could hear all cue differences; see below for details.

#### 3.2.3. Procedure

On each trial participants were presented with a sequence, then asked to click a button to indicate if it sounded more as if the beat was on every other note (“STRONG weak STRONG weak STRONG weak”) or every third note (“STRONG weak weak STRONG weak weak”). In total each participant completed 250 trials, ten blocks in each of which the 25 items were presented in random order. The experiment began with two practice trials.

#### 3.2.4. Analysis

For each participant logistic regression was conducted to examine the extent to which pitch versus duration influenced their categorization judgments. The outcome variable was the categorization decision for a given trial, with pitch (5 levels) and duration (5 levels) as predictors. The resulting coefficients were then normalized so that they summed to 1 as in Experiment 1. Comparisons among all three language groups are reported with the non-parametric Kruskal-Wallis H statistic. Comparisons of two language groups are reported with Mann-Whitney U tests (analogously to Experiment 1).

### 3.3. Results

Cue weights differed across language groups for musical beats (H(2) = 33.1, *p* < .001). Native Mandarin speaking participants had larger normalized pitch cue weights than both native English speakers (U = 1649, *p* < .001, A = 0.81) and Spanish speakers (U = 998, *p* < .001, A = 0.82), indicating that Chinese native speakers also relied on pitch to a greater degree when perceiving musical beats (Fig. 2a). English and Spanish speakers’ cue weights did not differ (U = 595, *p =* .89, A = 0.49). Figures 2b and 2c show how participants’ categorization responses changed as the pitch and duration levels of the stimuli were varied, respectively, collapsing across the other dimension; the categorization function for pitch was steeper for Mandarin speakers than for the other two groups, while the categorization function for duration was shallower for Mandarin speakers than for the other two groups. To investigate which stimuli were driving the group differences in categorization strategies, we used Mann-Whitney *U* tests to compare categorization responses to each of the 25 stimuli, using the Bonferroni method to correct for multiple comparisons. Group differences were almost exclusively confined to ambiguous stimuli in which pitch and duration information suggested conflicting interpretations (Figures 2d and 2e).

**Figure 2.**
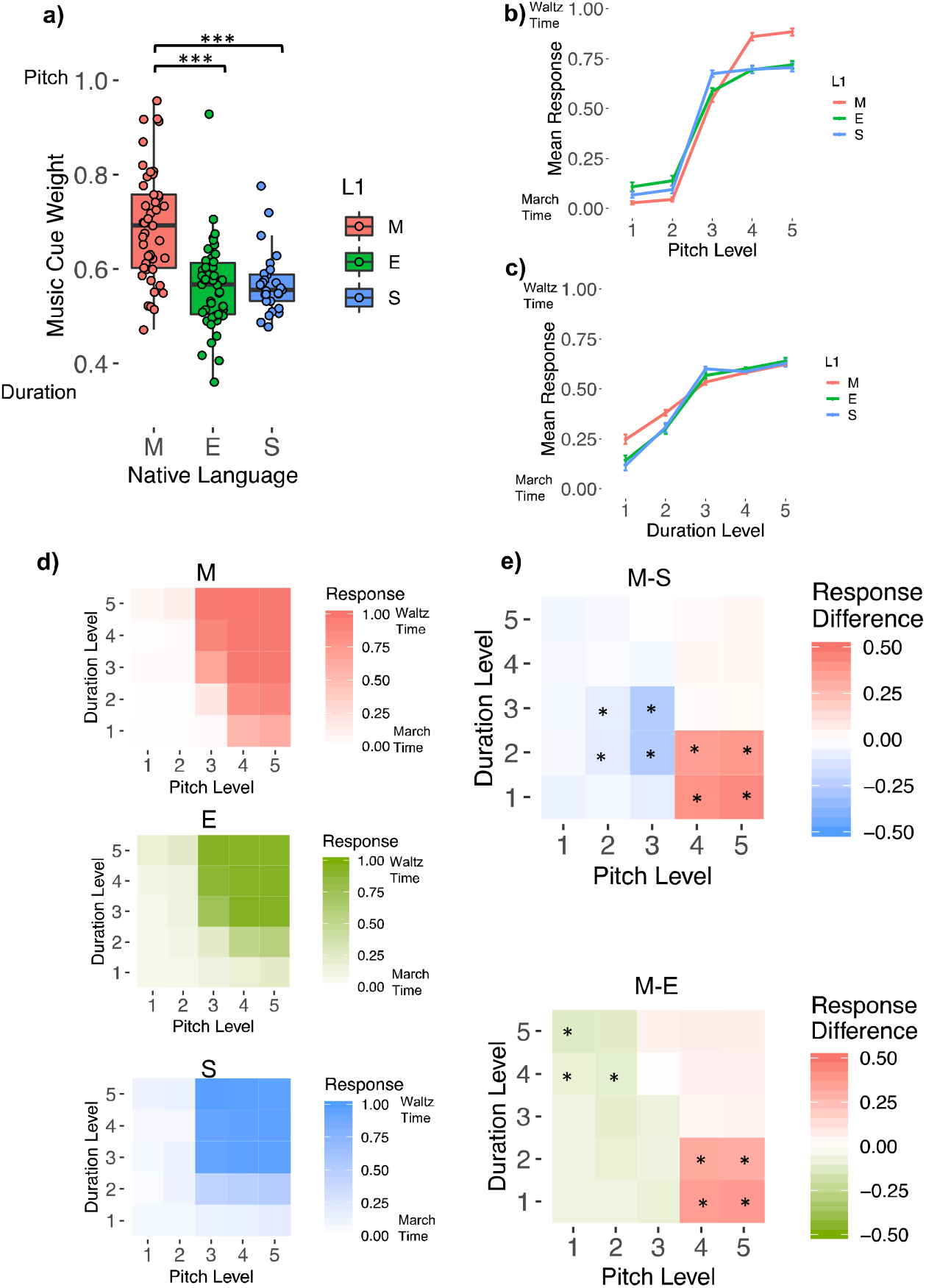
Mandarin speakers rely more on pitch and less on duration when categorizing features of music compared to English and Spanish speakers. **a)** Plots of normalized cue weights by language group for the music task. Greater values (approaching 1) indicate greater reliance on pitch, and lower values (approaching 0) reflect greater reliance on duration. **b)** Responses for levels of the pitch dimension (collapsed over duration) and **c)** for the duration dimension (collapsed over pitch). Error bars represent SEM. **d)** Plots of responses during the music task for each cell of the stimulus space, averaged across participants in each of the groups. Darker colors indicate more “late closure” responses. **e)** Responses made by the Mandarin group compared to the English and Spanish groups. Darker colors indicate relatively more “march time” responses made by the Mandarin (red), English (green) and Spanish (blue) groups. Asterisks indicate significant group differences in categorization responses, as tested using Mann-Whitney *U* tests with Bonferroni correction for multiple comparisons.

### 3.4. Discussion

Mandarin speakers’ judgments of beat locations reflected pitch information in the stimuli to a greater extent than the other two groups, suggesting they possess a perceptual strategy for music that matches their strategy for perceiving language —with a strong reliance on cues from pitch. This pitch-based strategy reflects their native experience with a tonal language. As we did for Experiment 1, we plotted each cell of the 5×5 stimulus space and found that the cue-incongruent corners of the space (especially the bottom right corner where pitch indicated a ‘triple meter’ but duration indicated a ‘tuple’ meter) showed the largest group differences. For those stimuli where pitch and duration conflicted, Mandarin speakers’ judgments diverged strongly from the other two groups as they relied on the pitch-based interpretation of the stimulus rather than the duration-based one. Moreover, again, we found that group differences were not limited to stimuli with more subtle pitch cues, but extended to stimuli with large, obvious pitch cues, suggesting that the group differences in categorization were not primarily driven by differences in sensitivity.

More generally, these results suggest that effects of perceptual experience on dimensional weighting are not limited to a particular perceptual task, or even a particular domain, but can extend across domains to potentially affect domain-general baseline perceptual strategies. These weighting shifts, therefore, may reflect modifications to auditory processing that occur relatively early (i.e. in regions of auditory cortex or the auditory midbrain sensitive to domain-general acoustic cues), rather than in down-stream domain-specific regions. For example, it has been previously shown that language speakers also show enhanced brainstem and cortical responses to pitch contours in verbal and non-verbal sounds (Chandrasekaran, Krishnan & Gandour 2007, Swaminathan, Krishnan & Gandour 2008, Chandrasekaran, Krishnan & Gandour 2009, Krishnan, Swaminathan & Gandour 2009, Bidelman, Gandour & Krishnan 2010, Krishnan, Gandour & Bidelman 2010, Bidelman et al. 2011, Krishnan, Suresh & Gandour 2019; but see Xu, Krishnan & Gandour 2006).

## 4. Experiment 3 - An attentional basis for domain-general perceptual strategies

### 4.1. Introduction

Experiments 1 and 2 found that Mandarin speakers relied more on pitch than non-tonal language speakers to perceive speech and music. What mechanism might explain effects of language experience on shifts in cue weighting that extend across domains? One possibility is that repeated task-relevance of a particular dimension leads it to become more salient, i.e. have a greater tendency to capture an individual’s attention, and that this increased salience leads the dimension to have greater influence during perceptual categorization (Gordon et al. 1993, Francis and Nusbaum 2002, Holt et al. 2018). In experiment 3 we tested one prediction of this account, which is that Mandarin speakers would have difficulty ignoring pitch and focusing on other information during a simple perceptual comparison task. We tested this hypothesis by repeatedly presenting to participants a two-word phrase, with the relative pitch height and amplitude of the two words varying orthogonally. On each trial participants judged which word they perceived to be either higher in pitch or greater in amplitude, while implicitly ignoring task-irrelevant changes from trial to trial along the other dimension.

Whereas in Experiments 1 and 2 there was no correct answer for a given trial (as participants simply indicated which interpretation of the stimulus they perceived), the stimuli of Experiment 3 did indeed have lower or high pitches and greater or less amplitude, and thus each trial had a correct answer. This allowed us to assess task performance as well as cue weights. Therefore, as in Experiments 1 and 2, we predicted that pitch variation in the stimuli should have a greater effect on judgments for Mandarin speakers than the non-tonal language groups, regardless of whether they are attending to pitch or amplitude. Furthermore, if pitch is more salient for Mandarin speakers, then they should perform better when tasked with attending to pitch and *worse* than the other two groups when tasked with attending to amplitude, because they will be distracted by the (irrelevant) pitch variation.

### 4.2. Methods

#### 4.2.1. Participants

The same participants took part as in Experiments 1 and 2.

#### 4.2.2. Stimuli

First, a recording was made of a voice actor reading aloud two different sentences: “Dave likes to STUDY music, but he doesn’t like to PLAY music” and “Dave likes to study MUSIC, but he doesn’t like to study HISTORY”. The fourth and fifth words of each recording—”study music”—were extracted; these recordings were, then, identical lexically but differed in the placement of contrastive focus (i.e. on “STUDY music” versus “study MUSIC”). The speech morphing software STRAIGHT (Kawahara & Irino 2005) was then used to morph these recordings onto one another so that the extent to which acoustic cues imply the existence of emphasis on one or the other word could be precisely controlled. (For more details see Jasmin, Dick, Tierney 2020c). Pitch and amplitude were then set to vary across four levels, from 0% (identical to the acoustic pattern for the recording with emphasis on the first word) to 33%, to 67%, to 100% (identical to the acoustic pattern for the recording with emphasis on the second word). The duration characteristics of each stimulus was identical, as the ‘time’ morphing parameter was always set to 50% (the average between the two original recordings).

#### 4.2.3. Procedure

For the “attend amplitude” condition, on each trial participants were presented with a single two-word phrase, then asked to say which word was louder. If the first word was louder, they clicked on a button marked “1”; if the second word was louder, they clicked on a button marked “2”. For the “attend pitch” condition, the procedure was the same, except that participants were asked to indicate which word was higher in pitch. Feedback was presented immediately after each trial in the form of a green check mark for correct responses and a red “X” for incorrect responses. Trial order was randomized. The “attend amplitude” condition was presented in its entirety first, followed by the “attend pitch” condition, to minimize task-switching effects. Each of the 16 stimuli was presented 3 times per condition, for a total of 48 trials per condition and 96 trials overall.

#### 4.2.4. Analysis

Portion correct was calculated separately for “attend amplitude” and “attend pitch” conditions. These values for the Mandarin, English and Spanish groups were compared with Mann-Whitney *U* tests. To investigate the effects of pitch and amplitude levels on responses, cue weights were calculated. For each participant logistic regression was conducted, with the outcome variable being the categorization decision for a given trial, with pitch (4 levels) and amplitude (4 levels) as predictors. The coefficients for pitch and amplitude from these regressions were normalized to sum to one using the formula from Section 2.2.4 and compared between groups with Mann-Whitney *U* tests. Comparisons among all three language groups are reported with the non-parametric Kruskal-Wallis H statistic.

### 4.3. Results - Dimension selective attention

#### 4.3.1. Attend pitch condition

The dimension selective attention task measured participants’ ability to attend to one dimension of speech (pitch or loudness) while simultaneously ignoring the other, independently varying dimension (loudness or pitch). The three groups differed in task performance during the attend-to-pitch condition (H(2) = 27.97, *p <* .001). Comparing the groups pair-wise revealed that, when asked to attend to pitch, Mandarin speakers had a greater proportion of correct judgments compared to speakers of English (U = 1513, *p* < .001, A = 0.74) and Spanish (U = 1022, *p* < .001, A = 0.84). Performance did not differ for the English and Spanish groups (U = 714.5, *p =* 0.21, A = 0.59).

The relative cue weights also differed across groups (H(2) = 22.17, *p <* .001). In line with this result, when asked to attend to pitch, pitch cues exhibited a stronger effect relative to amplitude cues on judgments for speakers of Mandarin than for speakers of English (U = 1462.5, *p* < .001, A = 0.72) or Spanish (U = 971, *p* < .001, A = 0.80; Fig. 3a). Pitch cue weights did not differ between the English and Spanish groups (U = 713.5, *p =* 0.22, A = 0.59). Figures 3b and 3c show how participants’ responses changed as the pitch and amplitude levels of the stimuli were varied, respectively, collapsing across the other dimension; the response function for pitch was steeper for Mandarin speakers than for the other two groups, while the response function for amplitude was shallower for Mandarin speakers than for the other two groups. To investigate which stimuli were driving the group differences in performance, we used Mann-Whitney *U* tests to compare responses to each of the 16 stimuli, using Bonferroni to correct for multiple comparisons, as in Experiments 1 and 2. Group differences were almost exclusively confined to ambiguous stimuli in which pitch and amplitude conflicted (Figures 3d and 3e).

**Figure 3.**
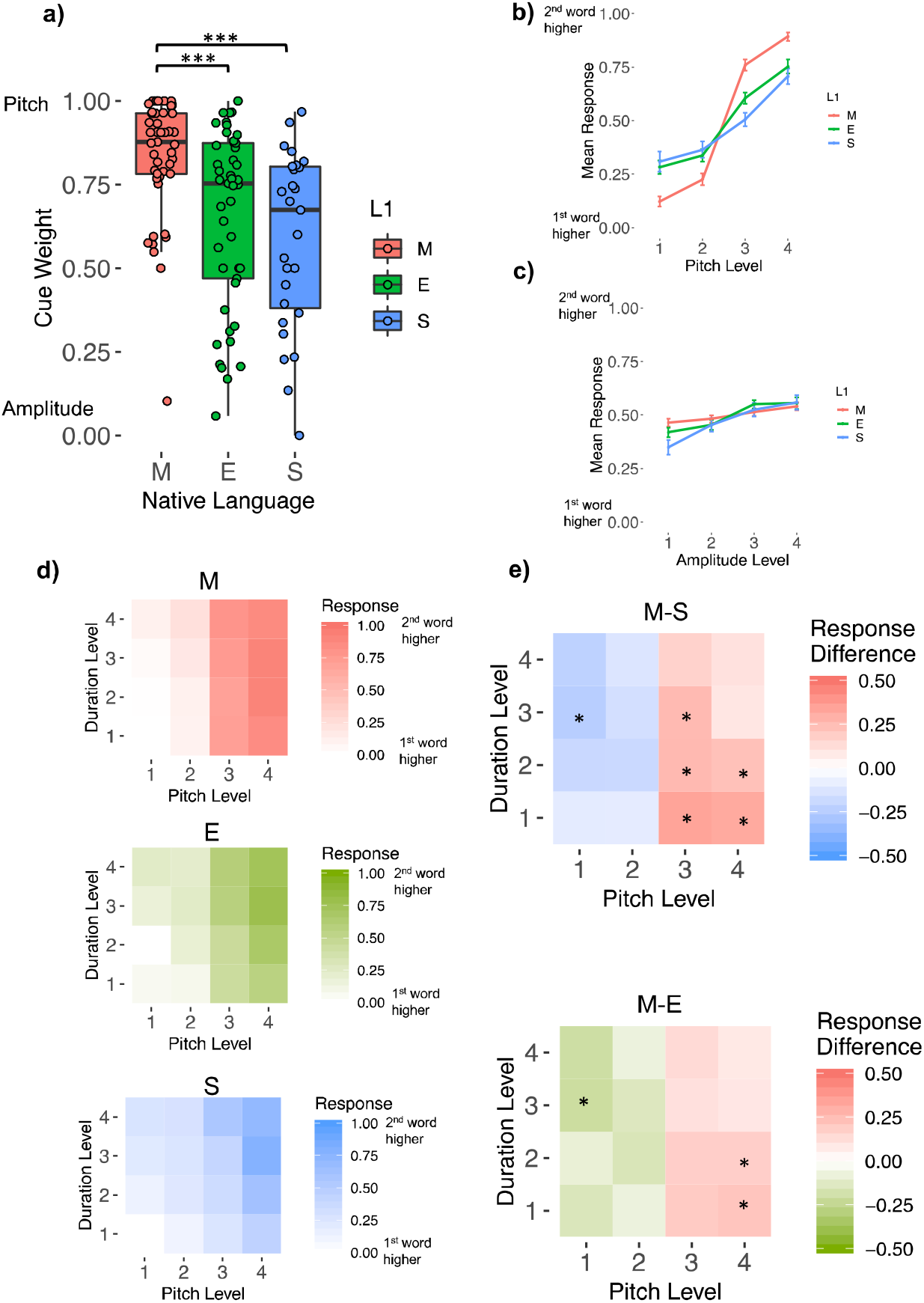
Mandarin speakers perform better when explicitly asked to attend to pitch information in speech, compared to English and Spanish speakers. **a)** Plots of normalized cue weights by language group for the attention-to-pitch task. Greater values (approaching 1) indicate greater influence of pitch on responses, and lower values (approaching 0) reflect greater influence of amplitude. **b)** Responses for levels of the pitch dimension (collapsed over amplitude) and **c)** for the amplitude dimension (collapsed over pitch). Error bars represent SEM. **d)** Plots of responses during the attention-to-pitch task for each cell of the stimulus space, averaged across participants in each of the groups. Darker colors indicate more “2nd word higher” responses. **e)** Responses made by the Mandarin group compared to the English and Spanish groups. Darker colors indicate relatively more “2nd word higher” responses made by the Mandarin (red), English (green), and Spanish (blue) groups. Asterisks indicate significant group differences in categorization responses, as tested using Mann-Whitney *U* tests with Bonferroni correction for multiple comparisons.

#### 4.3.2. Attend amplitude condition

In the attend-amplitude condition, the groups also differed in task performance (H(2) = 26.7, *p <* .001). Pair-wise group comparisons showed that when asked to attend to amplitude, Mandarin speakers showed lower performance than native speakers of English (U = 394, *p* < .001, A = 0.19) or Spanish (U = 328, *p =* .001, A = 0.27). English and Spanish speakers did not differ in performance (U = 710.5, *p =* 0.23, A = 0.58).

The three groups differed in the degree to which they relied on pitch or amplitude cues to make their judgments (normalized cue weights; (H(2) = 28.3, *p <* .001). Mandarin speakers exhibited significantly greater normalized pitch cue weights than speakers of English (U = 1646, *p* < .001, A = 0.81) and Spanish (U = 913.5, *p* < .001, A = 0.75; Fig. 4a). English and Spanish speakers’ pitch cue weights did not differ (U = 543, *p =* 0.46, A = 0.45). Figures 4b and 4c show how participants’ responses changed as the pitch and amplitude levels of the stimuli were varied, respectively, collapsing across the other dimension; the response function for pitch was steeper for Mandarin speakers than for the other two groups, while the response function for amplitude was shallower for Mandarin speakers than for the other two groups. To investigate which stimuli were driving the group differences in performance, we used Mann-Whitney *U* tests to compare responses to each of the 16 stimuli, using the Bonferroni method to correct for multiple comparisons. Group differences were exclusively confined to ambiguous stimuli in which pitch and amplitude conflicted (Figures 4d and 4e).

**Figure 4.**
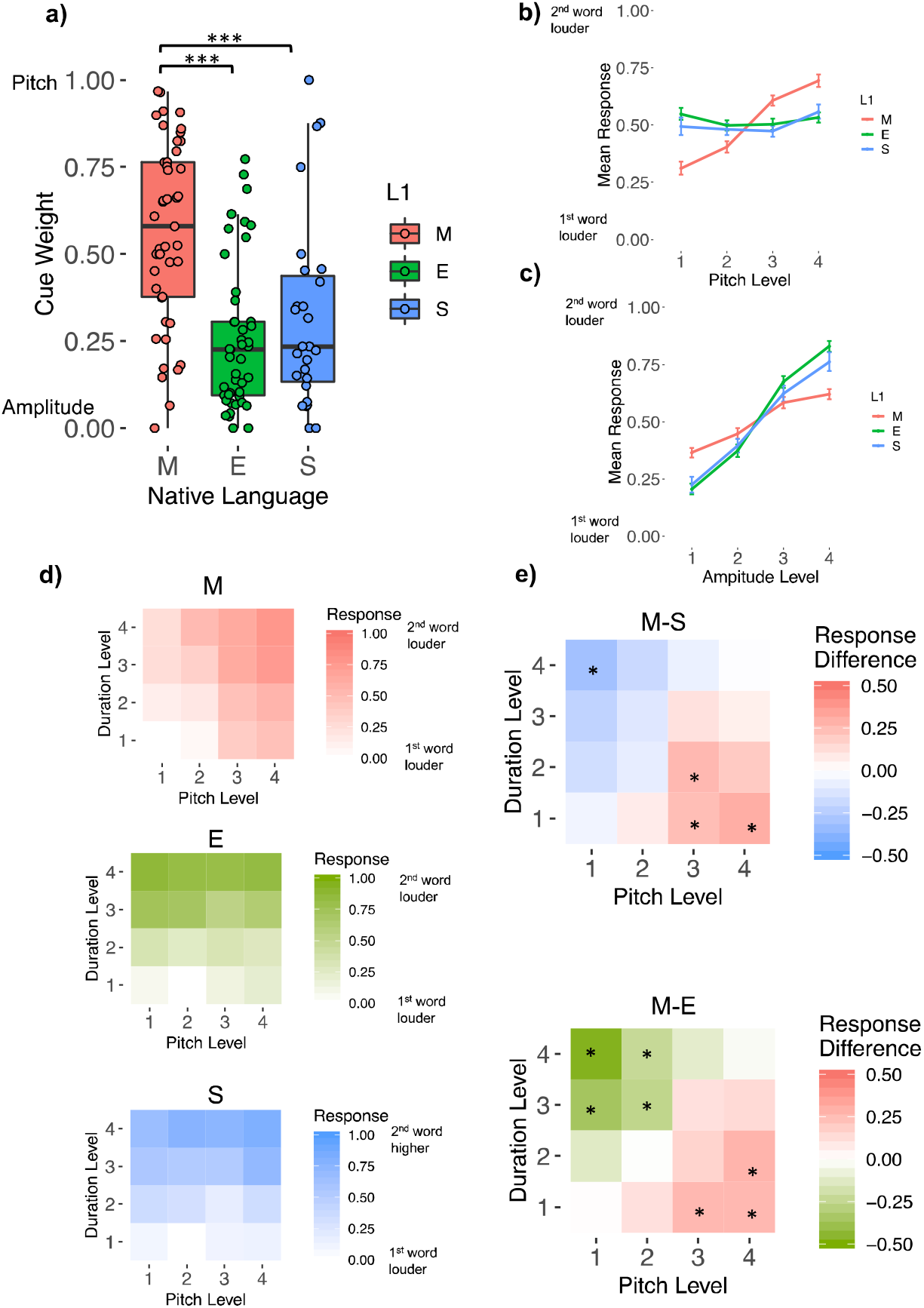
Mandarin speakers perform worse when explicitly asked to attend to amplitude information in speech, compared to English and Spanish speakers. **a)** Plots of normalized cue weights by language group for the attention-to-amplitude task. Greater values (approaching 1) indicate greater influence of pitch on responses, and lower values (approaching 0) reflect greater influence of amplitude. **b)** Responses for levels of the pitch dimension (collapsed over amplitude) and **c)** for the amplitude dimension (collapsed over pitch). Error bars represent SEM. **d)** Plots of responses during the attention-to-amplitude task for each cell of the stimulus space, averaged across participants in each of the groups. Darker colors indicate more “2nd word louder” responses. **e)** Responses made by the Mandarin group compared to the English and Spanish groups. Darker colors indicate relatively more “2nd word louder” responses made by the Mandarin (red), English (green) and Spanish (blue) groups. Asterisks indicate significant group differences in categorization responses, as tested using Mann-Whitney *U* tests with Bonferroni correction for multiple comparisons.

### 4.4. Discussion

Experiment 3 demonstrated a possible mechanism by which native language experience may shape auditory perception: via increased salience for an auditory dimension that is particularly important in one’s native language. Participants judged the amplitude and pitch height of a spoken stimulus while these two dimensions varied orthogonally. Compared to the non-tonal language groups (English and Spanish), speakers of Mandarin showed increased reliance on cues from pitch, even when explicitly directed to ignore pitch and attend to amplitude. This result suggests that Mandarin speakers experience increased pitch salience, and that this effect can be advantageous in certain circumstances but a hindrance in others: the Mandarin speakers were better at directing attention towards pitch and away from amplitude, but worse at directing attention towards amplitude and away from pitch. This inability to ignore pitch may be one factor leading Mandarin speakers to up-weight pitch cues during prosody perception in English, even when other cues are more important to native listeners (as in the perception of stress, in which vowel reduction is a more primary cue; Nguyễn et al. 2008, Wang, 2008, Zhang et al., 2008; Yu and Andruski, 2010; Zhang and Francis, 2010). One way to test this hypothesis would be to attempt to train native Mandarin speakers to explicitly attend to other dimensions in speech besides pitch, to see if this leads to shifts in cue weighting strategies during speech categorization, and whether this re-weighting extends to other speech perception tasks or to music perception.

The group differences in dimension-selective-attention performance also show that perceptual differences between speakers of Mandarin and non-tonal languages are not exclusively driven by increased pitch sensitivity, because a sensitivity account would not predict that Mandarin speakers would show impaired performance when asked to direct attention to amplitude. Indeed, we examined whether pitch sensitivity was correlated with “attend amplitude” performance across the 72 participants in the English and Spanish group, and there was no such correlation--rather, there was a trend in the opposite direction, with participants with the most precise pitch perception performing better on the amplitude task (Spearman rho = -.18, p=.12). These results suggest that perceptual sensitivity and salience are somewhat dissociable, and that it is pitch salience that is driving the group differences in selective attention performance in the tone-language versus non-tone-language speakers.

To further investigate whether group differences in sensitivity can explain our results, we ran a follow-up analysis in which we matched groups on pitch discrimination thresholds and re-ran all group comparisons across all three experiments.

## 5. Control Analyses in a Subsample of Participants Matched for Pitch Sensitivity

### 5.1. Introduction

As mentioned in the Introduction, several previous studies have demonstrated that tone language speakers tend to have finer pitch sensitivity (Bidelman et al., 2013; Giuliano et al., 2011; Hutka et al., 2015; Pfordresher and Brown, 2009; Zheng and Samuel, 2018). To investigate whether the results of our study are primarily driven by group differences in sensitivities, we measured pitch and duration sensitivity in each of our participants using pitch and duration discrimination tasks. The results of these tests were then used to exclude subjects, resulting in a reduced sample of participants whose pitch and duration thresholds were matched. The analyses in Experiments 1, 2, and 3 were then repeated in this new subset of participants in which sensitivity was not a confound.

### 5.2. Methods

#### 5.2.1. Stimuli

We created linear continua of 100 complex tones which varied on the basis of a single dimension, for both the pitch and duration tests. Stimuli were constructed from four-harmonic complex tones (equal amplitude across harmonics) with initial and final amplitude rise time of 15 ms (linear ramps) to avoid perception of clicks. For the pitch discrimination test, the baseline sound had a fundamental frequency of 330 Hz and the comparison sounds had fundamental frequencies which varied from 330.3 to 360 Hz, while the duration of the sounds was fixed at 500 ms. For the duration discrimination test, the baseline sound had a duration of 250 ms and the comparison sounds had durations which varied from 252.5 to 500 ms, while the fundamental frequency of the sounds was fixed at 330 Hz.

#### 5.2.2. Procedure

Psychophysical thresholds were recorded using a three-down one-up adaptive staircase procedure (Levitt, 1971). On each trial participants were presented with three sounds with a constant inter-stimulus-interval of 500 ms, with either the first sound or the last sound different from the other two. Participants were told to press either the “1” key or the “3” key on the keyboard to indicate which of the three sounds was different. No feedback was presented. The comparison stimulus level was initially set at step 50. The change in comparison stimulus level after each trial was initially set at 10 steps; in other words, the test became easier by 10 steps after every incorrect response and became more difficult by 10 steps after every third correct response. This step size changed to 5 steps after the first reversal, to 2 steps after the second reversal, and to 1 step after the third reversal and for the remainder of the test thereafter. Stimulus presentation continued until either 50 stimuli were presented or eight reversals were reached. Performance was calculated as the mean stimulus levels across all reversals from the second through the end of the test.

### 5.3. Results

Median duration thresholds did not differ across groups (M_M_ = 27 ms, IQR = 16-32; M_E_ = 25 ms, IQR = 16-42; M_S_ = 21 ms, IQR = 16-36; H(2) = 0.92, *p =* 0.63). Median pitch thresholds, however, did vary (M_M_ = 0.17 semitones, IQR = 0.10-0.26; M_E_ = 0.25 semitones, IQR = 0.15-0.40; M_S_ = 0.31 semitone, IQR = 0.22-0.53; H(2) = 11.9, *p* = .003). Mandarin speakers had lower thresholds than speakers of English (U = 744, *p* = .03, A = .37) and Spanish (U = 322.5, *p* < .001, A = 0.26). Spanish and English speakers’ thresholds did not differ (U = 473.5, *p* = .12). Importantly, all pitch thresholds were less than 1 semitone, ensuring that all pitch differences between stimuli were detectable by all participants. Similarly, all participants were capable of detecting a difference in duration of a factor of 2.

To ensure that the Mandarin speakers’ lower pitch detection thresholds were not driving the results of our experiments, we matched all three language groups for pitch thresholds by excluding the 10 Mandarin speakers with the lowest thresholds, 10 English speakers with the highest thresholds, and 10 Spanish speakers with the highest thresholds. In the resulting subset of participants, median pitch thresholds in the groups did not differ statistically (M_M_ = 0.20 semitones; M_E_ = 0.20 semitones; M_S_ = 0.24 semitones, Kruskal-Wallis H(2) = 0.98, *p* = 0.61, see Fig. 5).

**Figure 5:**
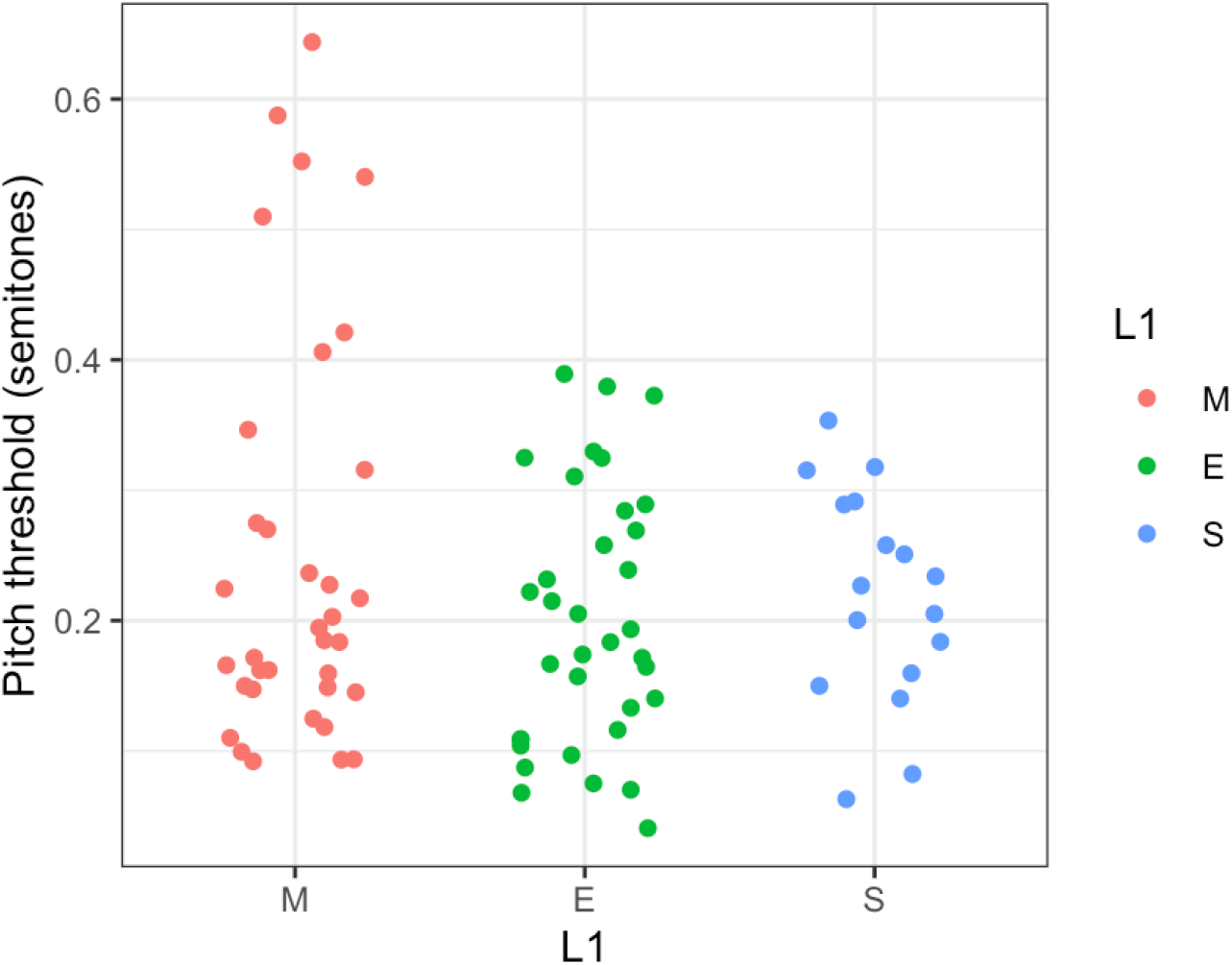
Pitch detection thresholds plotted by native language group, in a matched subset of participants.

Next, each of the statistical tests reported in the main text of the article was re-run in this matched subset of participants to determine if the inferences drawn in the article were upheld even when controlling for pitch sensitivity. We found that the results were not different from those in the full sample, such that all and only those test that were significant in the full sample were also significant in this subset, as described in Table 2. We therefore conclude that our results cannot be driven by a confound with pitch sensitivity. However, we wish to note that the sounds used in our pitch threshold task were static tones, whereas lexical tones often involve dynamic changes. Matching groups using a test of dynamic pitch discrimination may have provided a more stringent control.

**Table 2:** Results of the Main Analyses of the paper in the matched subset of participants

| *Experiment* | *Comparison* | *DV* | *Test Statistic* | *Test Statistic Value* | *p* |
| --- | --- | --- | --- | --- | --- |
| Expt 1 - Prosody | One-way ANOVA | Cue Weight | H(2) | 11.10 | **<.001** |
|  | M vs E | Cue Weight | U | 999 | **<.001** |
|  | M vs S | Cue Weight | U | 419 | **.02** |
|  | E vs S | Cue Weight | U | 200 | .06 |
| Expt 2 - Music | One-way ANOVA | Cue Weight | H(2) | 23.34 | **<.001** |
|  | M vs E | Cue Weight | U | 999 | **<.001** |
|  | M vs S | Cue Weight | U | 467 | **<.001** |
|  | E vs S | Cue Weight | U | 274 | .65 |
| Expt 3<br>Attend pitch | One-way ANOVA | Performance | H(2) | 11.97 | **.002** |
|  | M vs E | Performance | U | 829 | **.01** |
|  | M vs S | Performance | U | 465 | **.001** |
|  | E vs S | Performance | U | 339 | 0.42 |
|  | One-way ANOVA | Cue Weight | H(2) | 13.18 | **.001** |
|  | M vs E | Cue Weight | U | 813 | **.02** |
|  | M vs S | Cue Weight | U | 474 | **<.001** |
|  | E vs S | Cue Weight | U | 380 | .11 |
| Expt 3<br>Attend amplitude | One-way ANOVA | Performance | H(2) | 25.79 | **<.001** |
|  | M vs E | Performance | U | 185 | **<.001** |
|  | M vs S | Performance | U | 154 | **.005** |
|  | E vs S | Performance | U | 350 | .31 |
|  | One-way ANOVA | Cue Weight | H(2) | 23.6 | **<.001** |
|  | M vs E | Cue Weight | U | 1022 | **<.001** |
|  | M vs S | Cue Weight | U | 423 | **.015** |
|  | E vs S | Cue Weight | U | 220 | .133 |

## 6. General Discussion

In this study, we showed that native speakers of Mandarin, compared to native speakers of English and Spanish, have different perceptual strategies that are not limited to speech perception but extend to music perception as well. Mandarin speakers place more importance on pitch cues and less emphasis on durational cues compared to speakers of non-tonal languages, both when judging the locations of linguistic phrase boundaries and of musical beats. This suggests that Mandarin speakers’ extensive experience relying on pitch in the course of listening to speech has led to an increase in pitch weighting that is not limited to speech perception but extends to other domains. We also find that Mandarin speakers are better able to attend to pitch while ignoring amplitude changes in speech, but are impaired at attending to amplitude while ignoring pitch changes, suggesting that language experience can modify the extent to which perceptual dimensions are salient, i.e. liable to capture attention.

Recent computational models of cue weighting during speech perception suggest that dimensional weights are set for individual speech categorization tasks, via an assessment of the degree of separation of distributional peaks associated with discrete categories (Toscano and McMurray, 2010). Our results suggest, however, that cue weights may also be affected by more general baseline strategies, which reflect the relative past usefulness of perceptual dimensions across many different perceptual tasks. In other words, cue weights may reflect both local parameter setting (in response to distributions associated with a particular task) and global parameter setting (reflecting task-relevance across many different tasks). One way to test this idea would be to examine cue weighting in speech perception in musicians who speak non-tonal languages; unlike most speakers of non-tonal languages, musicians have needed to make extensive use of pitch information while perceiving and reproducing melodies, which could lead to a global up-weighting of pitch extending to speech perception.

One set of models which aim to explain the effects of language experience on dimensional weighting during categorization are dimension-selective-attention accounts, which suggest that up-weighted dimensions are more salient, tending to capture selective attention and therefore influence categorization to a greater extent (Gordon et al., 1993; Francis and Nusbaum, 2002; Holt et al., 2018). That language experience can have an effect on dimensional salience in speech is supported by the results of Experiment 3, which showed that Mandarin speakers continue to be greatly influenced by pitch information even when explicitly asked to direct their attention away from pitch and towards another dimension. Dimensional salience models of the effects of language experience on cue weighting are generally somewhat agnostic about whether a perceptual “dimension” is a domain-specific or domain-general phenomenon. Our results suggest that if language experience does lead to increased salience of perceptual dimensions, then this increased salience may not be limited to speech, suggesting that baseline dimensional salience patterns may cut across domain boundaries. One way to test this possibility would be to use fMRI to examine functional connectivity patterns during speech categorization, to see if dimensional weighting is linked to connectivity between executive control regions such as prefrontal cortex and early auditory regions dedicated to domain-general processing of specific auditory dimensions. We have published preliminary evidence that this is the case, showing that pitch weighting is tied to the degree of connectivity between left dorso-lateral prefrontal cortex and a right anterior superior temporal gyrus region which has previously been linked to non-verbal pitch perception (Jasmin et al., 2020b). Another way to test this possibility which could be addressed by future research would be to use EEG to examine neural entrainment to patterns of information across different dimensions in both verbal and non-verbal stimuli.

In our psychophysical testing we find more precise discrimination of the pitch of non-verbal tones in Mandarin speakers compared to speakers of non-tonal languages. However, we would argue for several reasons that the group difference in perceptual strategies we find cannot simply be a consequence of more precise pitch perception in the Mandarin speakers. First, we took care to ensure that across all three tasks, the size of the pitch differences between stimuli were greater than two semitones, well above participants’ discrimination thresholds. Second, increased pitch sensitivity cannot explain our finding that Mandarin speakers perform worse than non-tonal language speakers when asked to ignore pitch and attend to the amplitude of sounds. Lastly, we tested all effects reported in the paper in a subset of participants for whom pitch thresholds were equivalent in the three groups: all results persisted. Although this greater pitch weighting cannot simply be reduced to increased pitch sensitivity, there may be a relationship between the two advantages: decades of experience making use of pitch information during perceptual tasks may lead to low-level enhancements of the precision of pitch representations (consistent with the Reverse Hierarchy theory of perceptual learning, Ahissar and Hochstein, 2004).

One limitation of our study is that because our stimulus manipulations were large and therefore relatively obvious, participants may have been somewhat influenced by explicit awareness of the experimental design. This is a possibility that is hard to rule out, because it is difficult to know exactly what task strategies participants were using. However, our interpretation of our results is relatively unaffected by whether or not such explicit awareness took place. The results of Experiment 3 suggest that Mandarin speakers cannot help but up-weight pitch, even when they explicitly attempt not to do so. As a result, it is likely that the up-weighting of pitch in Mandarin speakers would be found regardless of whether or not they are explicitly aware of the fact that the experiment has been designed to assess their weighting of pitch versus another dimension.

In conclusion, here we show that native language experience shapes auditory perception in highly specific ways, not only in perception of other languages, but also for perception of other domains such as music. The results highlight a novel form of linguistic relativity: learning a language in which words are distinguished by a particular acoustic dimension affects perceptual strategies more generally.

## Author Contributions

All authors developed the study concept and contributed to the design. Testing and data collection were performed by H. Sun and A. Tierney. K Jasmin and A. Tierney performed the data analysis and interpretation. All authors drafted the manuscript and approved the final version of the manuscript for submission.

## Acknowledgments

This work was funded by a Leverhulme Trust Early Career Fellowship to KJ (ECF-2017-151), a Birkbeck Global Challenges Research Fund grant to HS, and a Research Project Grant from the Leverhulme Trust [grant number RPG-2019-107] to ATT.

We thank Aniruddh Patel for comments on an earlier version of this manuscript.

## References

Ahissar, M., & Hochstein, S. (2004). The reverse hierarchy theory of visual perceptual learning. Trends in Cognitive Sciences, 8(10), 457–464.

Anwyl-Irvine, A. L., Massonnié, J., Flitton, A., Kirkham, N., & Evershed, J. K. (2020). Gorilla in our midst: An online behavioral experiment builder. Behavior Research Methods, 52(1), 388–407.

Bailey, P., Dorman, M., & Summerfield, Q. (1977). Identification of sineLJwave analogues of CV syllables in speech and nonspeech modes. The Journal of the Acoustical Society of America, 61(S1), S66–S66.

Bent, T., Bradlow, A. R., & Wright, B. A. (2006). The influence of linguistic experience on the cognitive processing of pitch in speech and nonspeech sounds. Journal of Experimental Psychology: Human perception and performance, 32(1), 97.

Best, C. T., McRoberts, G. W., & Goodell, E. (2001). Discrimination of non-native consonant contrasts varying in perceptual assimilation to the listener’s native phonological system. The Journal of the Acoustical Society of America, 109(2), 775–794.

Bidelman G, Gandour J, Krishnan A (2010) Cross-domain effects of music and language experience on the representation of pitch in the auditory brainstem. Journal of Cognitive Neuroscience, 23, 425–434.

Bidelman G, Gandour J, Krishnan A (2011). Musicians and tone-language speakers share enhanced brainstem encoding but not perceptual benefits for musical pitch. Brain and Cognition, 77, 1–10.

Bidelman, G. M., Hutka, S., & Moreno, S. (2013). Tone language speakers and musicians share enhanced perceptual and cognitive abilities for musical pitch: evidence for bidirectionality between the domains of language and music. PloS one, 8(4), e60676.

Blicher, D. L., Diehl, R. L., & Cohen, L. B. (1990). Effects of syllable duration on the perception of the Mandarin Tone 2/Tone 3 distinction: Evidence of auditory enhancement. Journal of Phonetics, 18(1), 37–49.

Burns, E. M., & Sampat, K. S. (1980). A note on possible culture-bound effects in frequency discrimination. The Journal of the Acoustical Society of America, 68(6), 1886–1888.

Chandrasekaran, B., Krishnan, A., & Gandour, J. T. (2007). Mismatch negativity to pitch contours is influenced by language experience. Brain Research, 1128, 148–156.

Chandrasekaran, B., Krishnan, A., & Gandour, J. T. (2009). Relative influence of musical and linguistic experience on early cortical processing of pitch contours. Brain and Language, 108(1), 1–9.

Choi, W., Tong, X., & Samuel, A. G. (2019). Better than native: Tone language experience enhances English lexical stress discrimination in Cantonese-English bilingual listeners. Cognition, 189, 188–192.

Chrabaszcz, A., Winn, M., Lin, C. Y., & Idsardi, W. J. (2014). Acoustic cues to perception of word stress by English, Mandarin, and Russian speakers. Journal of Speech, Language, and Hearing Research, 57, 1468–1479. http://dx.doi.org/10.1044/2014_JSLHR-L-13-0279

Creel, S. C., Weng, M., Fu, G., Heyman, G. D., & Lee, K. (2018). Speaking a tone language enhances musical pitch perception in 3–5□year□olds. Developmental Science, 21(1), e12503.

Deroche, M. L., Lu, H. P., Kulkarni, A. M., Caldwell, M., Barrett, K. C., Peng, S. C., … & Chatterjee, M. (2019). A tonal-language benefit for pitch in normally-hearing and cochlear-implanted children. Scientific Reports, 9(1), 1–12.

Ellis, R. J., & Jones, M. R. (2009). The role of accent salience and joint accent structure in meter perception. Journal of Experimental Psychology: Human Perception and Performance, 35, 264–280. http://dx.doi.org/10.1037/a0013482

Ernst, M. O., & Banks, M. S. (2002). Humans integrate visual and haptic information in a statistically optimal fashion. Nature, 415(6870), 429–433.

Fear, B. D., Cutler, A., & Butterfield, S. (1995). The strong/weak syllable distinction in English. The Journal of the Acoustical Society of America, 97(3), 1893–1904.

Flege, J. E. (1995). Second language speech learning: Theory, findings, and problems. Speech Perception and Linguistic Experience: Issues in Cross-Language Research, 92, 233–277.

Francis, A. L., & Nusbaum, H. C. (2002). Selective attention and the acquisition of new phonetic categories. Journal of Experimental Psychology: Human perception and performance, 28(2), 349.

Fu, Q. J., & Zeng, F. G. (2000). Identification of temporal envelope cues in Chinese tone recognition. Asia Pacific Journal of Speech, Language and Hearing, 5(1), 45–57.

Giuliano, R. J., Pfordresher, P. Q., Stanley, E. M., Narayana, S., & Wicha, N. Y. (2011). Native experience with a tone language enhances pitch discrimination and the timing of neural responses to pitch change. Frontiers in Psychology, 2, 146.

Gordon, P. C., Eberhardt, J. L., & Rueckl, J. G. (1993). Attentional modulation of the phonetic significance of acoustic cues. Cognitive Psychology, 25(1), 1–42.

Hannon, E. E., Snyder, J. S., Eerola, T., & Krumhansl, C. L. (2004). The role of melodic and temporal cues in perceiving musical meter. Journal of Experimental Psychology: Human Perception and Performance, 30, 956–974. http://dx.doi.org/10.1037/0096-1523.30.5.956

Holt, L. L., Tierney, A. T., Guerra, G., Laffere, A., & Dick, F. (2018). Dimension-selective attention as a possible driver of dynamic, context-dependent re-weighting in speech processing. Hearing Research, 366, 50–64. http://dx.doi.org/10.1016/j.heares.2018.06.014

Hove, M. J., Sutherland, M. E., & Krumhansl, C. L. (2010). Ethnicity effects in relative pitch. Psychonomic Bulletin & Review, 17(3), 310–316.

Hutka, S., Bidelman, G. M., & Moreno, S. (2015). Pitch expertise is not created equal: Cross-domain effects of musicianship and tone language experience on neural and behavioural discrimination of speech and music. Neuropsychologia, 71, 52–63.

Idemaru, K., & Holt, L. L. (2011). Word recognition reflects dimension-based statistical learning. Journal of Experimental Psychology: Human Perception and Performance, 37, 1939–1956. http://dx.doi.org/10.1037/a0025641

Iverson, P., & Kuhl, P. K. (1994). Tests of the perceptual magnet effect for American English/r/and/l. The Journal of the Acoustical Society of America, 95(5), 2976–2976.

Iverson, P., Kuhl, P., Akahane-Yamada, R., Diesch, E., Tohkura, Y., Kettermann, A., & Siebert, C. (2003). A perceptual interference account of acquisition difficulties for non-native phonemes. Cognition, 87, B47–B57.

Jasmin, K., Dick, F., Holt, L. L., & Tierney, A. (2020a). Tailored Perception: Individuals’ Speech and Music Perception Strategies Fit Their Perceptual Abilities. Journal of Experimental Psychology: General. 149(5), 914 http://dx.doi.org/10.1037/xge0000688

Jasmin, K., Dick, F., Stewart, L., & Tierney, A. (2020b). Altered functional connectivity during speech perception in congenital amusia. eLife 9 (e53539)

Jasmin, K., Dick, F., & Tierney, A. T. (2020c). The Multidimensional Battery of Prosody Perception (MBOPP). Wellcome Open Research, 5(4), 4.

Kawahara, H., & Irino, T. (2005). Underlying principles of a high-quality speech manipulation system STRAIGHT and its application to speech segregation. In P. Divenyi (Ed.), Speech separation by humans and machines (pp. 167–180). Boston, MA: Kluwer Academic Publishers. http://dx.doi.org/10.1007/0-387-22794-6_11

Krishnan, A., Gandour, J. T., & Bidelman, G. M. (2010). The effects of tone language experience on pitch processing in the brainstem. Journal of Neurolinguistics, 23(1), 81–95.

Krishnan, A., Suresh, C. H., & Gandour, J. T. (2019). Tone language experience-dependent advantage in pitch representation in brainstem and auditory cortex is maintained under reverberation. Hearing Research, 377, 61–71.

Krishnan, A., Swaminathan, J., & Gandour, J. T. (2009). Experience-dependent enhancement of linguistic pitch representation in the brainstem is not specific to a speech context. Journal of Cognitive Neuroscience, 21(6), 1092–1105.

Levitt, H. C. C. H. (1971). Transformed upLJdown methods in psychoacoustics. The Journal of the Acoustical Society of America, 49(2B), 467–477.

Lin, H. B., & Repp, B. H. (1989). Cues to the perception of Taiwanese tones. Language and Speech, 32(1), 25–44.

Lisker, L. (1986). “Voicing” in English: A catalogue of acoustic features signaling /b/ versus /p/ in trochees. Language and Speech, 29(1), 3–11.

Liu, S., & Samuel, A. G. (2004). Perception of Mandarin lexical tones when F0 information is neutralized. Language and Speech, 47(2), 109–138.

Lutfi, R. A., & Liu, C. J. (2007). Individual differences in source identification from synthesized impact sounds. The Journal of the Acoustical Society of America, 122(2), 1017–1028.

Massaro, D. W., & Cohen, M. M. (1977). Voice onset time and fundamental frequency as cues to the /zi/-/si/ distinction. Perception & Psychophysics, 22(4), 373–382.

Mattys, S. L. (2000). The perception of primary and secondary stress in English. Perception & Psychophysics, 62, 253–265. http://dx.doi.org/10.3758/BF03205547

Nguyễn, T. A. T., Ingram, C. J., & Pensalfini, J. R. (2008). Prosodic transfer in Vietnamese acquisition of English contrastive stress patterns. Journal of Phonetics, 36(1), 158–190.

Palmer, C., & Krumhansl, C. L. (1987). Independent temporal and pitch structures in determination of musical phrases. Journal of Experimental Psychology: Human Perception and Performance, 13, 116–126. http://dx.doi.org/10.1037/0096-1523.13.1.116

Peretz, I., Nguyen, S., & Cummings, S. (2011). Tone language fluency impairs pitch discrimination. Frontiers in Psychology, 2, 145.

Pfordresher, P. Q., & Brown, S. (2009). Enhanced production and perception of musical pitch in tone language speakers. Attention, Perception, & Psychophysics, 71(6), 1385–1398.

Plag, I., Kunter, G., & Schramm, M. (2011). Acoustic correlates of primary and secondary stress in North American English. Journal of Phonetics, 39(3), 362–374.

Prince, J. B. (2014). Pitch structure, but not selective attention, affects accent weightings in metrical grouping. Journal of Experimental Psychology: Human Perception and Performance, 40, 2073–2090. http://dx.doi.org/10.1037/a0037730

Stagray, J. R., & Downs, D. (1993). DIFFERENTIAL SENSITIVITY FOR FREQUENCY AMONG SPEAKERS OF A TONE AND A NONTONE LANGUAGE/使用 声调语言和非声调语 言为母语的人对声音频率的分辨能力. Journal of Chinese Linguistics, 143-163.

Streeter, L. A. (1978). Acoustic determinants of phrase boundary perception. Journal of the Acoustical Society of America, 64, 1582–1592. http://dx.doi.org/10.1121/1.382142

Swaminathan, J., Krishnan, A., & Gandour, J. T. (2008). Pitch encoding in speech and nonspeech contexts in the human auditory brainstem. Neuroreport (11), 1163.

Tierney, A. T., Russo, F. A., & Patel, A. D. (2011). The motor origins of human and avian song structure. Proceedings of the National Academy of Sciences of the United States of America, 108, 15510–15515. http://dx.doi.org/10.1073/pnas.1103882108

Toscano, J. C., & McMurray, B. (2010). Cue integration with categories: Weighting acoustic cues in speech using unsupervised learning and distributional statistics. Cognitive Science, 34(3), 434–464.

Wang, Q. (2008). Perception of English stress by Mandarin Chinese learners of English: An acoustic study (Doctoral dissertation). University of Victoria. Retrieved from https://venus.library.uvic.ca:8443/bitstream/handle/1828/1282/Dissertation_2008Dec09.pdf?sequence=1&isAllowed=y

Whalen, D. H., & Xu, Y. (1992). Information for Mandarin tones in the amplitude contour and in brief segments. Phonetica, 49(1), 25–47.

Winter, B. (2014). Spoken language achieves robustness and evolvability by exploiting degeneracy and neutrality. BioEssays, 36(10), 960–967.

Wong, P. C., Ciocca, V., Chan, A. H., Ha, L. Y., Tan, L. H., & Peretz, I. (2012). Effects of culture on musical pitch perception. PloS one, 7(4), e33424.

Xu, Y., Krishnan, A., & Gandour, J. T. (2006). Specificity of experience-dependent pitch representation in the brainstem. Neuroreport, 17(15), 1601–1605.

Yu, V.Y. & Andruski, J. E. (2010). A cross-language study of perception of lexical stress in English. Journal of Psycholinguistic Research, 39(4), 323–344.

Zhang, X. (2012). A Comparison of Cue-Weighting in the Perception of Prosodic Phrase Boundaries in English and Chinese (Doctoral dissertation). University of Michigan. Retrieved from https://deepblue.lib.umich.edu/bitstream/handle/2027.42/96107/zxt_1.pdf?sequence=1

Zhang, Y., & Francis, A. (2010). The weighting of vowel quality in native and non-native listeners’ perception of English lexical stress. Journal of Phonetics, 38(2), 260–271.

Zhang, Y., Nissen, S. L., & Francis, A. L. (2008). Acoustic characteristics of English lexical stress produced by native Mandarin speakers. The Journal of the Acoustical Society of America, 123(6), 4498–4513.

